# MetaChrome: An Open-Source, User-Friendly Tool for Automated Metaphase Chromosome Analysis

**DOI:** 10.1101/2025.09.02.673813

**Authors:** Md Abdul Kader Sagar, Yamini Dalal, Gianluca Pegoraro, Ganesan Arunkumar

## Abstract

DNA Fluorescence In Situ Hybridization (FISH) is an essential technique to study chromosome biology and genetics, enabling precise visualization of specific genomic loci to study structural abnormalities, gene mapping, and chromosomal rearrangements. High-Throughput Imaging (HTI) can automate the analysis of DNA-FISH chromosome images, but the accurate and automated segmentation of mitotic chromosomes and simultaneous colocalization of FISH signals remains a challenge. While several commercial automated karyotyping tools partially solve these issues, open-source software that effectively combines robust chromosome segmentation with comprehensive colocalization analysis capabilities remains necessary. To address this unmet need, we developed MetaChrome, an open-source software platform built around a graphical user interface and explicitly designed for automated metaphase chromosome analysis. MetaChrome leverages fine-tuned deep learning models to automate metaphase chromosome segmentation, together with colocalization analysis of chromosome-specific FISH probes and immunofluorescent-labeled proteins. Importantly, MetaChrome achieves enhanced segmentation accuracy compared to traditional image processing methods by adopting a Cellpose segmentation model fine-tuned with manually annotated metaphase chromosome datasets. The fine-tuned model ensures precise assignment of DNA-FISH spots to individual chromosomes in an automated manner. This facilitates rapid identification of chromosomal abnormalities, reduces human error, and advances high-throughput chromosome analysis workflows, addressing a key bottleneck in chromosome biology research.

## 1. Introduction

Metaphase chromosomes are prominent structures observed during cell division, specifically in the metaphase stage of mitosis or meiosis [1]. At this stage, chromosomes are highly condensed and aligned at the cell equatorial plane [2], making them easier to visualize under a microscope. Somatic human cells contain 46 chromosomes, organized into 23 pairs: 22 pairs of autosomes and one pair of sex chromosomes [3]. Chromosomes contain the cell’s genetic material, and, in humans, alterations in their morphology are diagnostic of diseases such as cancer. Therefore, analysis of metaphase chromosomes is now essential for a host of applications, including genetic research, cancer diagnostics, and the identification of chromosomal abnormalities during development [4]. Accurate karyotyping, which involves the classification and examination of chromosomes from images of metaphase spreads, is crucial for diagnosing genetic disorders and understanding the genetic basis of diseases. Karyotyping enables the detection of structural and numerical chromosome alterations, informing clinical decisions and therapeutic interventions. Consequently, precise and efficient karyotyping plays a fundamental role in personalized medicine and improved patient outcomes [5,6].

Metaphase chromosome segmentation is a specialized area within clinical and basic science research and represents a critical step in chromosome analysis [7]. A relevant example of a subtle, yet significant, chromosomal aberration is the mislocalization of the centromeric protein A (CENP-A) to the 8q24 chromosomal arm, a region frequently amplified in various cancers [8]. Understanding the clinical relevance of this event requires accurately measuring its occurrence across many images of metaphase spreads. Unfortunately, though, manual segmentation and classification of chromosomes are time-consuming processes that require a high level of expertise, leading to potential errors and inconsistencies. This specific need highlights a broader challenge in the karyotyping field: the accurate segmentation of chromosomes from metaphase spreads images, which is essential for precisely characterizing chromosomal markers. Therefore, developing a reliable, semi-automated analysis pipeline to quantify this specific molecular mislocalization is essential for achieving the scale and accuracy needed for robust clinical research.

Recent advancements in High-Throughput Imaging (HTI) [9], telomere-to-telomere sequencing [10], and high-resolution DNA Fluorescence In Situ Hybridization (FISH) have enabled researchers to examine metaphase chromosomes in greater detail. These techniques facilitate the identification of microdeletions, translocations, amplifications, and inversions that were previously detected using nucleic acid staining methods such as G-banding [11–14].

Several automated karyotyping tools, such as IdeoKar [15], ChromosomeJ [16], ChromaWizard [17], and KaryoXpert [18] excel at segmenting images of metaphase chromosomes stained with trypsin- or pepsin-based banding techniques under brightfield microscopy. However, they face significant limitations analyzing, segmenting chromosomes, and colocalizing labeled DNA-FISH probes and target proteins labeled for immunofluorescent detection. Although commercial software solutions, including CytoVision and MetaSystems Ikaros, are available, their high costs make them inaccessible for individual lab setups and basic science laboratories, particularly those with limited resources and personnel. This limitation is especially critical for chromosome biologists and clinicians analyzing specific genomic loci associated with copy number variations, translocations, or epigenetic modifications. Such research often demands precise localization and quantification of protein and RNA signals on metaphase chromosomes, as well as the ability to perform multiplexed fluorescence analysis [19,20].

Over the past decade, deep learning-based instance segmentation models have emerged as state-of-the-art solutions for complex biomedical image segmentation tasks [21–23]. However, applying these models to chromosome segmentation poses specific and significant challenges. A primary difficulty is separating overlapping and clustered chromosomes, a common problem in object instance segmentation that is compounded by high variability in chromosome morphology. Furthermore, there is a scarcity of accurately labeled public datasets, as their manual generation is exceptionally labor-intensive and requires expert knowledge. Finally, variations in metaphase image quality and complexity—arising from differences in experimental conditions, fluorescence probes, and imaging systems—not only introduce artifacts but also complicate the consistent evaluation and comparison of segmentation performance across different studies.

To overcome these challenges, we developed MetaChrome, an open-source, cross-platform computational tool specifically designed to streamline metaphase chromosome segmentation and analysis from fluorescence microscopy data in a semi-automated manner. The software takes advantage of a Cellpose cell segmentation model [24,25], fine-tuned using an expert-annotated fluorescence microscopy image dataset of metaphase chromosomes to adapt the model to mitotic chromosome segmentation. Next, MetaChrome includes a user-friendly graphical interface built upon the Napari visualization platform [26], allowing researchers to intuitively interact and curate segmented chromosome regions of interest (ROI), localize DNA-FISH spots, and quantify fluorescence signals effectively. By simplifying and enhancing these steps, such as automated chromosome segmentation and counting, FISH signal spot detection on segmented chromosomes, and signal intensity measurements for co-localizing signals from different imaging channels, MetaChrome greatly accelerates the process of chromosomal abnormality detection, improves analytical reproducibility, and supports efficient, high-throughput chromosome analysis workflows. For example, relative to manual analysis, we have found that analyzing colocalization of two fluorescence signals from different channels across 40 metaphase spreads typically takes about 1-2 hours. The same analysis in MetaChrome will typically take less than 2 minutes using GPU computing, or less than 10 minutes using CPU computing. Therefore, we believe this tool will enable researchers across disciplines to perform mitotic chromosome analysis with greater speed, accuracy, and reproducibility.

## 2. Methods

### 2.1 Metaphase chromosome preparation

SW480 human colon cancer cells (ATCC, CCL-228), grown and maintained in our lab [8,27,28], were cultured in RPMI 1640 medium (Catalog no. 11875093, Gibco, USA) with glutathione, 10% fetal bovine serum (Catalog no. S11050H, Atlanta Biologicals, USA), and 1x penicillin-streptomycin solution (Catalog no. 15140122, Gibco, USA) at a temperature of 37°C and 5% CO2 in a humidified incubator. Cells were grown in a T175 flask until 60% confluency, and colcemid (Catalog no. 10295892001, Millipore Sigma, USA) was added 8 hours before the end of harvest time. The cells were harvested through trypsinization (Catalog no. 25200056, Thermo Fisher Scientific, USA), washed with 10 ml of 1X PBS, then suspended in 6 ml of hypotonic solution (0.075 M KCl), and incubated in a water bath at 37°C for 20 min. They were then fixed using a freshly prepared cold fixative solution (methanol and glacial acetic acid in a 3:1 v/v ratio). After centrifugation at 1500 rpm for 5 minutes at room temperature, cells were suspended in 4 ml of fixative and incubated for 15 minutes at room temperature. Finally, they were centrifuged again, resuspended in 400 μl of the fixative solution, and stored at 4°C until slide preparation.

### 2.2 Immunofluorescence staining and in situ hybridization

We utilized Immunofluorescence and DNA-fluorescence in situ hybridization (FISH) protocols as previously described in detail [28]. Metaphase chromosomes were dropped onto glass slides and allowed to air-dry. The slides were incubated in TEEN buffer, which consists of 1 mM triethanolamine-HCl (pH 8.5), 0.2 mM Na-EDTA, and 25 mM NaCl, and continued for immunofluorescence staining. A blocking step was performed using 0.1% Triton X-100 and 0.1% BSA in the TEEN buffer. Primary antibody incubation was conducted in KB buffer, composed of 10 mM Tris-HCl (pH 7.7), 150 mM NaCl, and 0.1% BSA, for 4 hours. After washing with KB buffer, the slides were incubated with secondary antibodies conjugated to AlexaFluor dyes (ThermoFisher) for 45 minutes at room temperature. The slides are washed with KB buffer before crosslinking using 10% Formaldehyde in KB buffer for 10 minutes. The slides are then washed with water and processed for DNA FISH hybridization.

For DNA FISH, slides were equilibrated in 2X SSC buffer at room temperature and digested with Pepsin (Catalog no. P6887, Sigma-Aldrich, USA) in 2X SSC at 10 mg/ml (Catalog no. 10108057001, Millipore Sigma, USA) at 37°C. The slides were washed in 2X SSC and dehydrated through an ethanol series (70%, 95%, and 100%). Denaturation was performed in 70% formamide in 2X SSC at 80°C, followed by another dehydration step. The DNA FISH probe targeting the 8q24 locus tagged with the FITC fluorophore (Empire Genomics, USA) was then added to the slides in a 10 μl volume of hybridization buffer (50% formamide, 2x SSC, 0.5mM EDTA, and 10% dextran sulfate). The slides were covered with a coverslip, sealed with rubber cement, and incubated in a humidified chamber at 37°C overnight. After washes with 2X SSC containing 50% formamide and 0.2X SSC, the slides were air-dried and mounted using a DAPI-containing mounting solution in the dark.

### 2.3 Imaging

Fluorescence imaging was performed using a Delta Vision™ Elite RT microscopy system (GE Healthcare, USA) equipped with a CoolSNAP CCD camera (Photometrics, USA) mounted on an inverted IX-70 microscope (Olympus, USA). DNA-FISH and immunofluorescence slides were imaged using a 60× oil immersion objective lens (NA 1.42). Optical filter turrets were configured for DAPI (441–459 nm), DNA-FISH targeting the 8q24 locus (501–549 nm; probe peak at 525 nm), and immunofluorescence detection of CENP-A tagged with Alexa Fluor 647 (662–696 nm). Exposure times were 250 ms for DAPI, 500 ms to 1.5 s for DNA-FISH, depending on slide age and photobleaching, and 800 ms for CENP-C immunofluorescence. Images were acquired at 1021 × 1024 pixels, 96 dpi resolution, and 16-bit depth, then deconvolved, processed, and exported as uncompressed TIFF files using softWoRx software (Cytiva, USA).

### 2.4 Cellpose fine-tuning and inference

The Cellpose model was fine-tuned on a dataset of hand-annotated metaphase chromosomes, which was split into 70% training, 15% validation, and 15% test subsets using the Python module scikit-learn [29] ‘train_test_split’ function. The model was initialized with pre-trained Cellpose-cyto model weights (Pachitariu and Stringer 2022) for fine-tuning, and training was performed with a batch size of 4, learning rate of 0.001, and the Adam optimizer for 100 epochs. We first generated an expanded train set on disk with a Python script that reads each TIFF/mask pair, applies random 0°/90°/180°/270° rotations, horizontal/vertical flips, and brightness scaling, and writes out five augmented variants per image. During training, we then invoked the Cellpose command line interface CLI with the --augment flag, which tiles each input into overlapping patches and applies random flips, 90° rotations, and brightness/contrast adjustments on-the-fly. Training was stopped early whenever the validation loss failed to improve for five consecutive epochs. The code and instructions to run MetaChrome can be found here https://github.com/CBIIT/MetaChromeTool2024. The associated documentation for the project can be found here https://napari-chromosome-analysis.readthedocs.io/.

### 2.5 Napari graphical user interface development

To make this powerful segmentation model accessible to researchers with minimal or no programming skills, the MetaChrome graphical user interface (GUI) was developed as a self-contained plugin for the Napari multi-dimensional image viewer. The main interface is constructed as a QWidget using PyQt5, which is then registered with Napari to appear as a dockable widget. This widget organizes the entire workflow into distinct, collapsible sections for each stage of the analysis, such as image loading, segmentation, and spot detection.

The tool uses Napari’s native layer-based system to manage and display data. For instance, after segmentation, the resulting masks are added to the viewer as a new Labels layer, which overlays the original DAPI image layer. User interactions for manual curation are directly tied to Napari’s event-driven architecture. Manual edits, such as splitting/merging/deleting chromosomes, and deleted spots are captured using a dedicated Shapes layer, allowing the user to draw lines or regions that are then used by the backend code to modify the segmentation mask. This integration of custom processing logic with Napari’s inherent visualization and interaction capabilities creates a seamless workflow for the end-user. MetaChrome can be installed directly from the Python Package Index (PyPI) by running the command pip install metachrome. Additional installation details and usage instructions are available at the official PyPI page: https://pypi.org/project/metachrome/1.0.0/.

### 2.6 Hardware requirements

MetaChrome is designed to be broadly accessible, running on any platform that supports both the Conda package and environment management software, and the Napari viewer. The primary requirement is a system capable of managing a Conda environment, which handles the installation of all necessary dependencies, including PyQt5 for the graphical user interface. For reliable operation, a modern multi-core processor (e.g., Intel Core i7 or equivalent) and a minimum of 16 GB of system RAM are also recommended, especially when handling large, multi-channel images. While a dedicated GPU is not required, the type of user hardware will directly impact analysis speed. The software can run entirely on a standard multi-core CPU, although deep learning-based segmentation will be considerably slower than with an NVIDIA GPU with CUDA support. For example, using 1024 × 1024, 16-bit, 2–3-channel TIFF images (∼6 MB each), Cellpose inference on an 8-core CPU processes approximately 0.2–0.3 images/s. On an NVIDIA RTX 3060, throughput increases to 2–3 images/s (∼10x faster), on an RTX 4090 to 5–6 images/s (∼20x faster), and on an NVIDIA A100 (40 GB) to 8–10 images/s (∼35–40x faster). Given the main bottleneck for the analysis is the segmentation, using a standard GPU like RTX 3060, a 40-image dataset takes less than a minute for the whole pipeline to be finished, assuming user intervention is not needed for modifying segmentation.

### 2.7 Data export and analysis

To ensure consistency across different ASO-mediated knockdown timepoints, MetaChrome batch image analysis was run with the same parameters across different image datasets. Following analysis in MetaChrome, quantitative single object data—including spot coordinates, fluorescence intensities, and colocalization status—were exported as CSV files for each experimental condition. These datasets were then read, structured, and managed using the pandas library [30]. The resulting individual data files were then consolidated into a single master table containing all measurements and corresponding experimental metadata for comprehensive analysis.

The primary measurement of this analysis was the 8q24/CENP-C colocalization ratio, which was calculated for each experimental condition. A DNA-FISH spot (marking the 8q24 locus) and a CENP-C immunofluorescence signal were defined as colocalized if the distance between them was less than a user-defined threshold, set to 12 pixels based on optimization through visual inspection. The final ratio represents the percentage of chromosomes meeting this proximity criterion. For direct comparison of protein levels, CENP-C fluorescence intensities were normalized to the average intensity of the scramble controls and expressed as a fold-change relative to this baseline to account for experimental variability.

To assess temporal changes within treatment conditions, statistical comparisons between the 72h and 96h timepoints were performed using independent samples t-tests. The resulting data was visualized using grouped box plots created with matplotlib, which show the complete distribution of normalized intensities.

A comparative analysis of CENP-C intensity at the centromere was performed across three timepoints (72h, 96h, and 120h) for both the ASO Scramble control and the ASO PCAT2 knockdown cells. For this comparison, the intensity data were first log2-transformed. We then generated grouped box plots to visualize the intensity trends side-by-side for the control and knockdown groups over the experimental time course. All the Python code used to generate the plots in Figs. 3 and 4 can be found in the GitHub repository for the code (https://github.com/CBIIT/metachromePaperCode).

### 2.8 Quantification of Model Performance

To quantitatively assess the performance of our segmentation methods, we utilized these metrics:

*Intersection over Union (IoU)*: measures the overlap between the predicted segmentation mask and the ground truth mask. It is calculated as the area of intersection divided by the area of union of the two masks. Higher IoU values indicate better spatial agreement between the predicted and actual segmentation.

*Precision*: measures the proportion of correctly identified positive chromosome segments (true positives) among all chromosomes identified as positive by the model (true positives + false positives). It indicates the accuracy of the positive predictions.

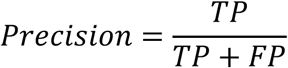

*Recall*: measures the proportion of correctly identified positive chromosome segments (true positives) among all actual positive segments in the ground truth. It indicates the model’s ability to find all relevant instances.

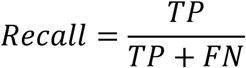

*F1-score*: harmonic mean of precision and recall. It provides a balanced measure of a model’s accuracy, particularly useful when there is an uneven class distribution.

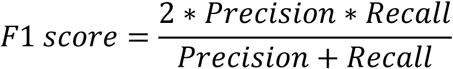

## 3. Results

### 3.1 Automated Workflow of MetaChrome for Metaphase Chromosome Analysis

The analysis of metaphase chromosomes is key to studying genetic instability and chromosomal abnormalities linked to cancer and genetic disorders. In particular, imaging of DNA-FISH and IF stained metaphase spreads can be used, among other applications, to quantify changes in protein abundance at specific genomic sites [28], recombination events [31], measure telomere length [32], and microsatellite amplification [33]. For example, in certain cancer cells, the centromeric protein CENP-A is ectopically deposited at non-centromeric loci like the 8q24 chromosomal arm, an event that requires precise measurement to understand its clinical relevance [8,27,28,34,35]. A recent study demonstrated that RNA Polymerase II (RNAPII) occupancy at histone genes is strongly correlated with tumor aggressiveness across various cancer types. Consequently, assessing RNAPII occupancy using DNA-FISH targeted to histone genes combined with co-immunofluorescence for RNAPII offers a rapid and cost-effective alternative to sequencing and bioinformatics, potentially enhancing clinical management of aggressive tumors [36]. However, traditional analysis of such events relies on manual visual inspection of fluorescence microscopy images, a process that is laborious, subjective, and error-prone due to user fatigue. With modern microscopy experiments becoming increasingly complex and data-rich, there is a pressing need for automated computational methods that enhance the efficiency, accuracy, and reproducibility of mitotic chromosome analysis.

To this end, we created a computational workflow to segment metaphase chromosomes and localize spot-like IF and/or DNA-FISH signals in fluorescence microscopy images. The analysis workflow, which we incorporated into our software tool named MetaChrome, can be broadly divided into four main stages (Fig. 1). As the first step, users begin by loading and visualizing fluorescence microscopy images captured from multiple channels if MetaChrome is run in interactive mode to setup the analysis parameters for spot finding (Fig. 1A). Typically, these include a DNA-binding fluorescent dye (like DAPI or Hoechst) for metaphase chromosomes, DNA-FISH probes highlighting specific genomic regions (in this study, the 8q24 locus), and IF markers for protein of interest (here, centromeric protein CENP-C). MetaChrome then directs the loaded images into its analytical pipeline for chromosome segmentation and subsequent spot analyses. The second analysis workflow step involves automated mitotic chromosome segmentation using a Cellpose deep learning model on the DAPI image (Fig. 1B). In interactive mode, the user can correct segmentation errors using a point and click interface in the Metachrome graphical user interface (GUI). Users have the flexibility to delete incorrectly segmented chromosomes, merge those mistakenly separated, or split chromosomes that are incorrectly merged. Manual corrections are entirely optional and only required when automated segmentation does not meet specific requirements. In parallel, Metachrome uses a local contrast-based peak finding algorithm to localize spot-like signals in the IF and DNA-FISH channels, respectively (Fig. 1C). While the detection process is automated, the software provides options to manually correct spot detection, allowing users to eliminate any false-positive signals or imaging artifacts. Finally, MetaChrome identifies chromosomes exhibiting co-localized DNA-FISH and protein signals (Fig. 1D). This feature is crucial for studies investigating the co-localization of proteins and their intensity at specific genomic loci. Finally, MetaChrome precisely measures fluorescence intensities at these defined spots, providing detailed quantitative data. All data measured for the segmented chromosomes, identified spots, and intensity measurements can easily be exported in CSV format, facilitating further statistical and comparative analyses (Fig. 1D).

**Figure 1.**
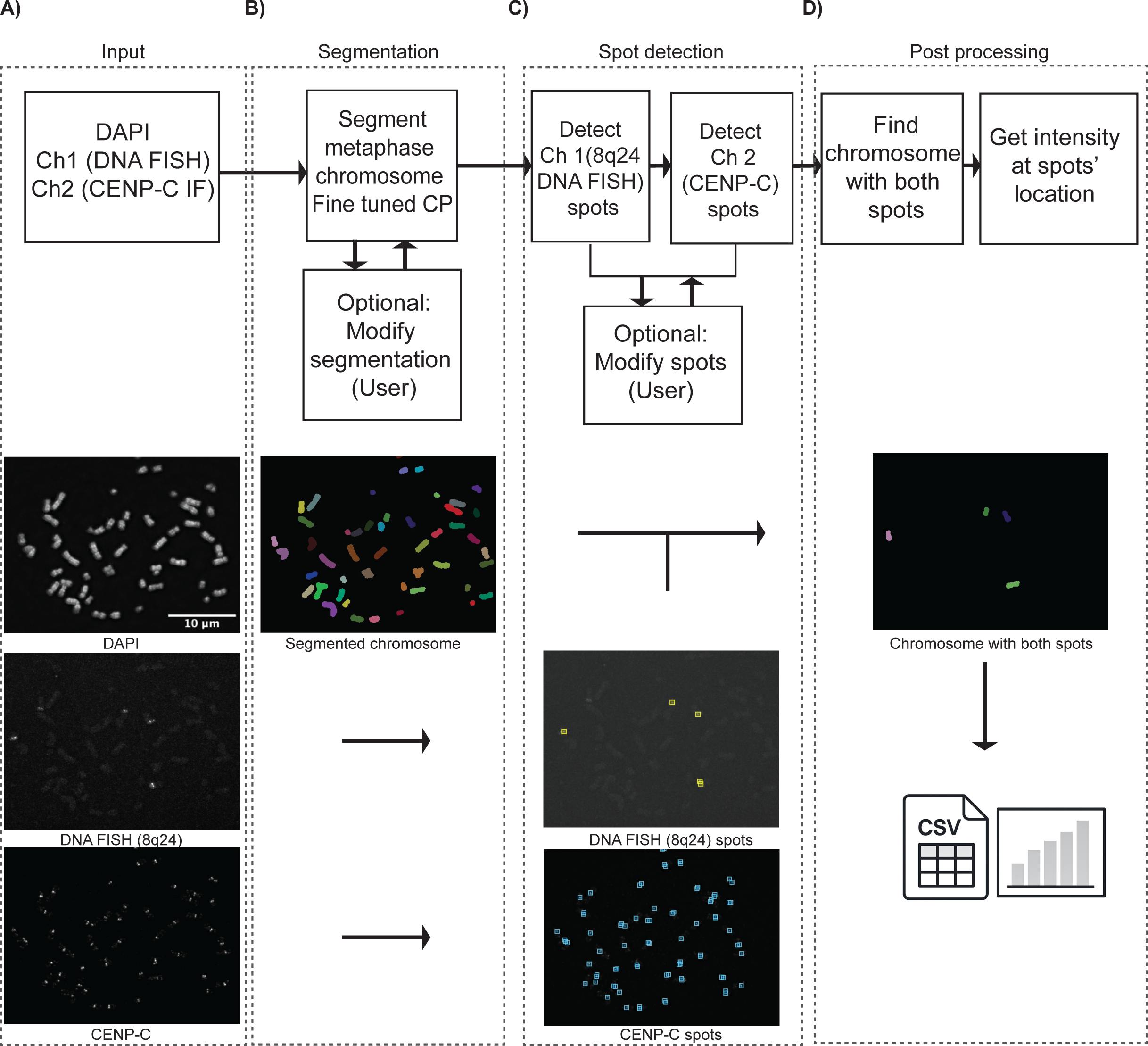
MetaChrome Workflow for Automated Metaphase Chromosome Analysis. The workflow consists of four key stages. A) Initially, raw fluorescence microscopy images from multiple channels (Channel 1: DAPI, Channel 2: DNA FISH (8q24) and CENP-C) are loaded into MetaChrome for segmentation and analysis. B) Metaphase chromosomes are automatically segmented using a retrained Cellpose model, providing accurate chromosome boundaries. Users have the option to manually adjust segmentation results if necessary. C) DNA FISH spots corresponding to the 8q24 loci (Channel 1) and CENP-C protein localization (Channel 2) are detected automatically, with optional manual correction to refine accuracy. (iv) Chromosomes containing both DNA FISH and protein spots are identified, enabling quantitative intensity measurements at spot locations. The final data, including chromosome identities and corresponding quantitative metrics, can be exported as CSV files for downstream analysis.

### 3.2 MetaChrome Software Architecture and User Interface

MetaChrome is designed with a modular, user-centric architecture that leverages widely-used Python computational libraries and frameworks, including PyTorch [37], scikit-image [38], SciPy [39], NumPy [40], and Napari [26] (Figure 2A). The Napari framework provides a graphical user interface that allows users without programming experience to visualize images, modify spot detection parameters, select analysis workflow options, and launch the analysis in batch mode on multiple images. To this end, the GUI (Figure 2B) is organized into several interactive panels that guide the user through a logical analysis workflow. It allows for streamlined data navigation and interactive visualization, which substantially enhances usability and the efficiency of reviewing results. Users begin by loading images from specified folders, with channels identified by unique string identifiers like fluorescence excitation wavelengths (e.g., DAPI at 435 nm, 8q24 at 525 nm, and CENP-C at 679 nm in our use case, Figure 2Bi). All operations and images are managed as distinct layers that can be toggled for clear visualization using Napari’s built-in functions and additional buttons added for MetaChrome. The remaining panels in Figure 2B (ii–vii) highlight the various GUI functionalities outlined in the figure caption.

**Figure 2.**
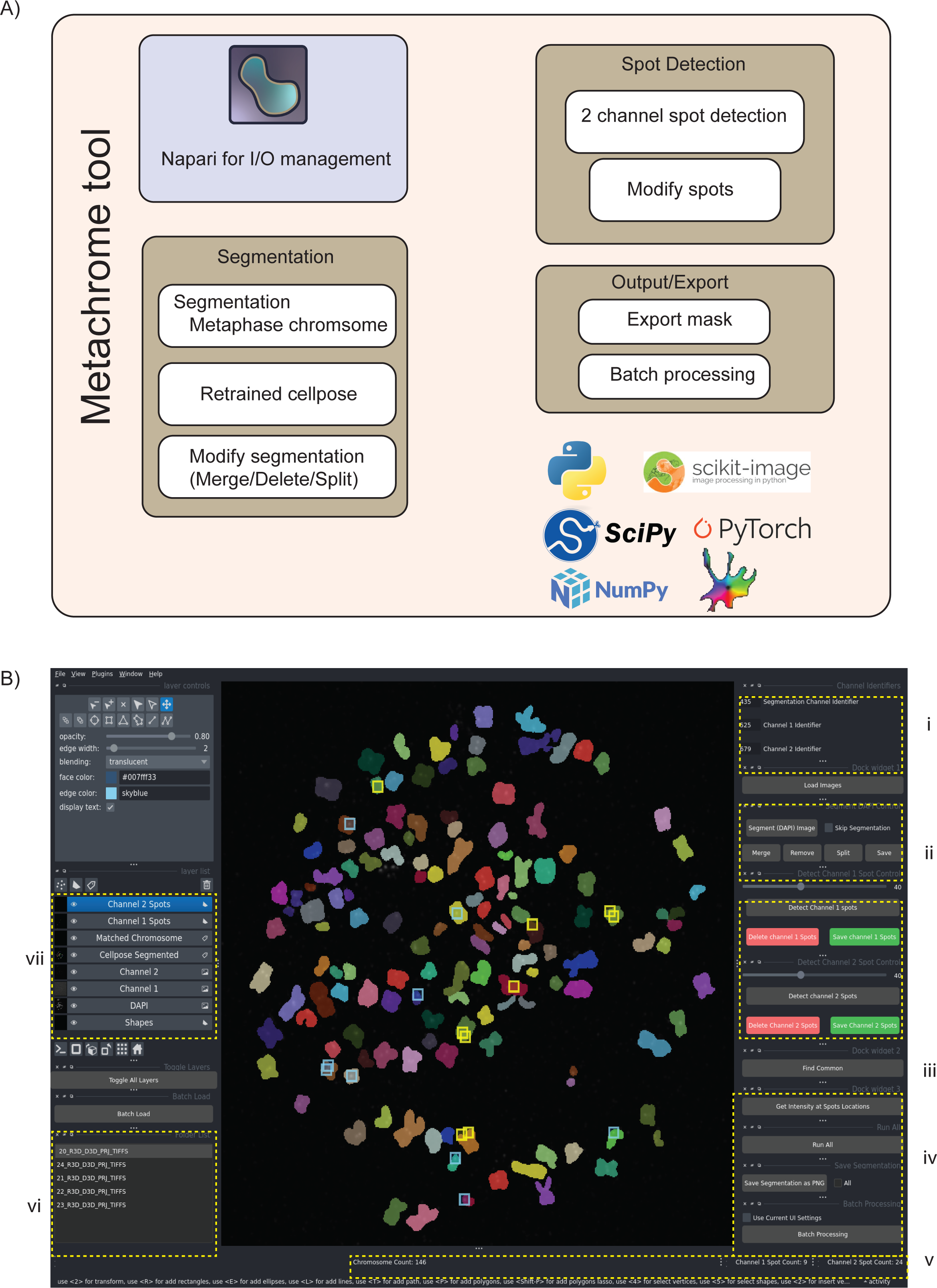
MetaChrome Software Architecture and User Interface. A) Schematic diagram outlining the software components of MetaChrome, highlighting its integration of widely used Python libraries, including PyTorch, scikit-image, SciPy, NumPy, and the Napari visualization framework. MetaChrome provides automated segmentation through a retrained Cellpose model, enabling manual editing of segmented chromosomes (merging, splitting, and deleting). It also performs two-channel spot detection and supports batch processing and export functionalities. B) Screenshot of the MetaChrome graphical user interface, built using Napari, organized into multiple interactive sections: i) Image loading and channel identification; ii) Chromosome segmentation and editing controls; iii) Spot detection controls, including manual adjustments; iv) Intensity calculation and batch processing options; v) Real-time chromosome and spot counts; (vi) List of loaded image files for quick access and batch operations; and vii) Display of various Napari layers with different steps of processed images.

The operational workflow starts with automated chromosome segmentation from the DAPI channel via the retrained Cellpose model. The software integrates a fine-tuned Cellpose segmentation model optimized specifically for accurate metaphase chromosome segmentation, along with user-friendly editing tools for manual refinement, such as merging, deleting, or splitting segmented objects. If users modify a segmentation, they can save it for reproducibility or export it for future retraining purposes. The MetaChrome workflow also allows users to integrate their own custom models that have been retrained on specific datasets. Users can also integrate their own custom object segmentation models retrained on specific datasets into the MetaChrome workflow. Similar to the chromosome segmentation step, users can manually remove incorrectly identified spots to refine the analysis. A key function in this section automatically identifies chromosomes that contain co-localized spots from multiple channels, directly linking different molecular signals.

For quantitative analysis, the software calculates fluorescence intensity values at the locations of the detected spots. The interface provides a real-time display of the number of chromosomes and spots, which updates with each modification, offering immediate feedback to the user. To facilitate high-throughput studies, a batch processing mode allows the settings from an interactive session to be applied to all open files.

Besides the workflow described above, which incorporates segmentation, MetaChrome also includes a “spots-only” analysis mode, where users can quantify intensity purely based on detected spots, without requiring chromosome segmentation. This mode will enhance the scoring of DNA-FISH signals from marker chromosomes, extra-chromosomal DNAs, and chromothripsis. In this mode, intensity calculations utilize all detected spots across both fluorescence channels independently of DAPI-based segmentation, and the user can still modify the detected spots using a user-defined threshold and use MetaChrome’s toolset to manually delete unwanted spots. Moreover, MetaChrome’s chromosome segmentation and counting feature alone can be used for chromosome counting and to detect aneuploidy and macro-chromosomal features. Overall, this semi-automated image analysis workflow streamlines and accelerates metaphase chromosome analysis, increases reproducibility, and minimizes the need for manual intervention for mitotic chromosome analysis.

### 3.3 Comparative Evaluation of Mitotic Chromosomes Segmentation Performance

To develop a robust and effective mitotic chromosome segmentation method, we systematically evaluated several segmentation approaches. Using representative metaphase chromosome images stained with DAPI and ground truth segmentation annotations (Fig. 3A), we qualitatively assessed performance through visual inspection, and quantitatively using precision, recall, and F1-scores across varying intersection-over-union (IoU) thresholds—a common approach for quantifying segmentation of biological objects in fluorescence microscopy images [41]. Please see the Materials and Methods section for details. Our first test of a classical watershed algorithm proved insufficient for this task (Fig. 3B). It displayed considerable segmentation errors in distinguishing overlapping chromosomes and performed poorly from a quantitative standpoint (F1 score = 0.12 at 0.7 IoU threshold, Fig. 3C).

**Figure 3.**
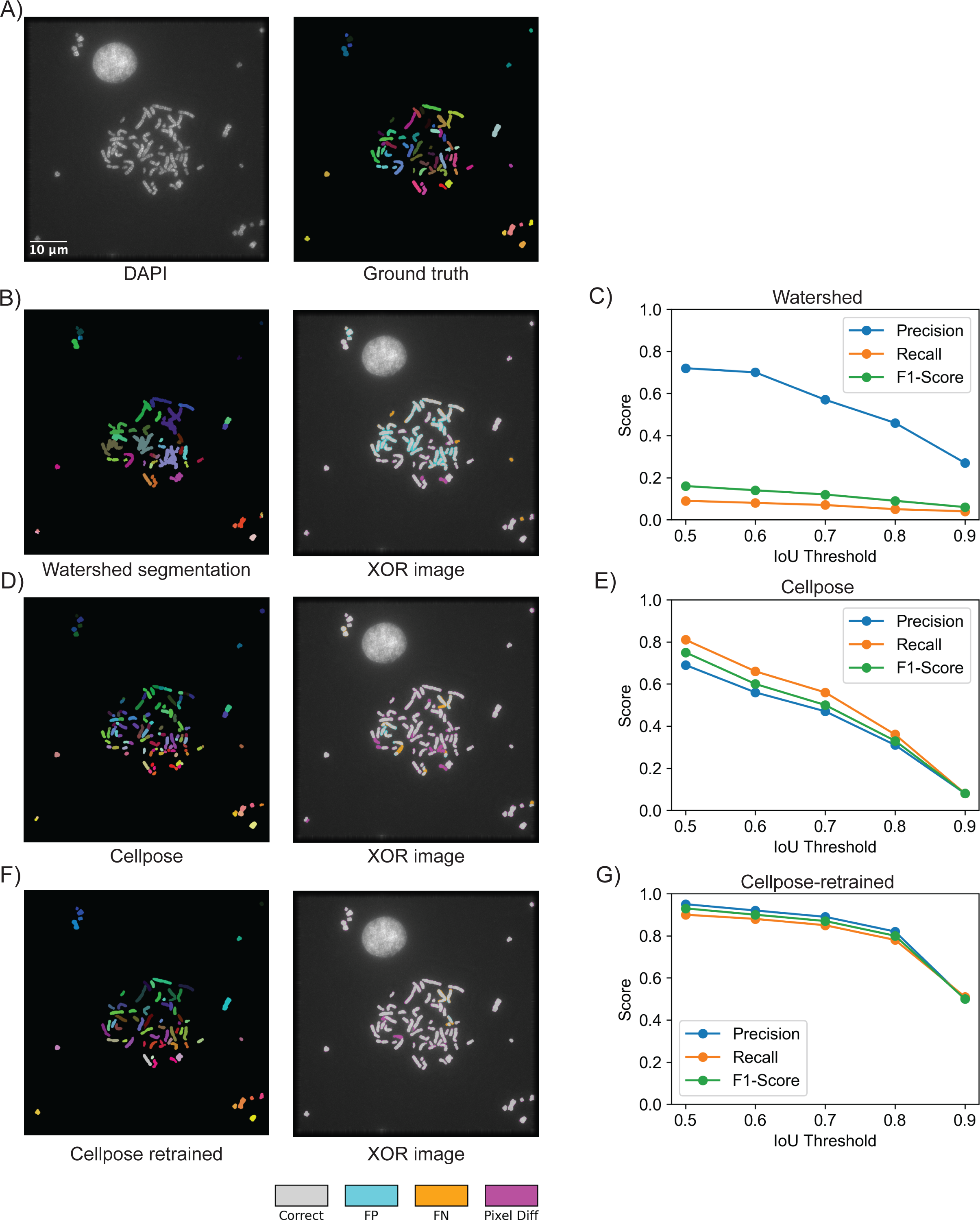
Comparative Evaluation of Chromosome Segmentation Methods. (A) Representative metaphase chromosome image stained with DAPI (left) alongside its corresponding manually annotated segmentation mask (ground truth, right). (B) Example segmentation result using the watershed segmentation method (left), with differences highlighted by XOR overlay (right), illustrating segmentation inaccuracies. (C) Quantitative evaluation of watershed segmentation performance using precision, recall, and F1-score across varying IoU thresholds, demonstrating significant performance limitations. (D) Segmentation outcome using the original Cellpose model (left), with corresponding XOR image (right) highlighting segmentation errors. (E) Performance metrics for the original Cellpose model show improved but still limited accuracy at higher IoU thresholds. (F) Enhanced segmentation using the retrained Cellpose model (left), with XOR overlay (right), demonstrating significantly improved segmentation accuracy. (G) Quantitative analysis confirms superior segmentation performance (precision, recall, F1-score) achieved by the retrained Cellpose model, maintaining high accuracy even at stringent IoU thresholds. The evaluation was performed on 7 FOVs not used for training or validation. In XOR overlays, gray = correct overlap, blue = false positives, orange = false negatives, magenta = severe instance mismatch.

Given these limitations, we next evaluated the default, pre-trained Cellpose deep learning model. This model performed better compared to the watershed algorithm (F1 score 0.51 at IoU 0.7, Fig. 3D), but still showed significant inaccuracies, particularly at higher IoU thresholds (F1 score 0.33 at IoU 0.8, Fig. 3E). Therefore, for our final approach, we adopted a transfer learning strategy suggested by the Cellpose model authors [25] and fine-tuned the pre-trained model using our own manually annotated dataset of mitotic chromosomes (See Materials and Methods for details). Compared to the original Cellpose model, the Cellpose model fine-tuned on mitotic chromosomes images demonstrated improved segmentation accuracy (Fig. 3F), effectively overcoming the challenges posed by the non-regular shape of chromosomes and closely adjacent chromosomes. Quantitative assessment (Fig. 3G) confirmed that our fine-tuned model produced high F1 scores both at standard IoU threshold (0.87 at IoU 0.7) and, more importantly, also at high IoU thresholds (0.84 at IoU 0.8). Collectively, these results indicate that MetaChrome’s fine-tuned Cellpose model significantly outperforms conventional segmentation approaches, providing researchers with a reliable tool for precise metaphase chromosome segmentation necessary for advanced quantitative analyses. Furthermore, we also provide users with guidelines on how to additionally fine-tune the Cellpose model for use cases where segmentation of metaphase chromosomes with different morphological features is needed, and on how to integrate the fine-tuned models into an existing MetaChrome pipeline (https://napari-chromosome-analysis.readthedocs.io/).

### 3.4 MetaChrome Automates the Analysis of Multiplexed Fluorescence Microscopy Images of Mitotic Chromosomes

As an example of a practical MetaChrome application, and to validate its use as a semi-automated image analysis pipeline for mitotic chromosomes analysis, we used Metachrome to analyze a previously published dataset [28] of images of mitotic chromosomes stained with IF and DNA-FISH.

The goal of the experiments that generated this imaging dataset was to investigate the role of the chromosome 8q24-derived long non-coding RNA PCAT2 in maintaining the mislocalization of CENP-A, which is normally localized at the centromeres, at the same non-centromeric locus in certain cancer cells [8,27,28]. The results of those experiments showed that the ASO-mediated knockdown of PCAT2, which physically associates with and facilitates ectopic deposition of CENP-A at its transcribing locus, resulted in significant depletion of CENP-A from the 8q24 locus in SW480 cells [28]. This was confirmed by imaging the cells stained in IF for CENP-C, a CENP-A binding protein [28]. Here, we used MetaChrome to reanalyze images of SW480 mitotic chromosomes from cells where PCAT2 was knocked down for 72, 96, and 120 hours, and that were stained with a CENP-C antibody and a DNA-FISH probe against the 8q24 locus (Figure 4A). Figure 4B shows the spot colocalization ratio of CENP-C at the 8q24 locus. The normalized intensity shown in Figure 4C was the CENP-C signal intensity data at the 8q24 locus obtained from biological replicates. To measure changes in CENP-C localization at 8q24, MetaChrome was used to analyze a total of 6,312 mitotic chromosomes in a total of 101 images. MetaChrome post-processing analysis revealed a reduction in CENP-C intensity at the 8q24 locus 96 hours post PCAT2 knockdown. This observation is consistent with our visual scoring of CENP-C colocalization at the 8q24 locus on metaphase chromosomes that was previously published [28]. In addition to CENP-C fluorescence intensity, MetaChrome could also reproduce, in an automated manner, the significant reduction in the percentage of CENP-C foci colocalized at the 8q24 genomic site at 72, 96, and 120 hours post-transfection of PCAT2 ASO when compared to the scramble control [28] (Fig. 4B). The ability of MetaChrome to accurately quantify these changes in CENP-C levels further underscores its effectiveness as a robust tool for studying alterations in chromosomal protein localization. Overall, the results of this re-analysis show that MetaChrome can accurately perform metaphase chromosome segmentation and protein/DNA colocalization (both scoring intensities and colocalizing spot counts) in an automated and unbiased manner.

**Figure 4.**
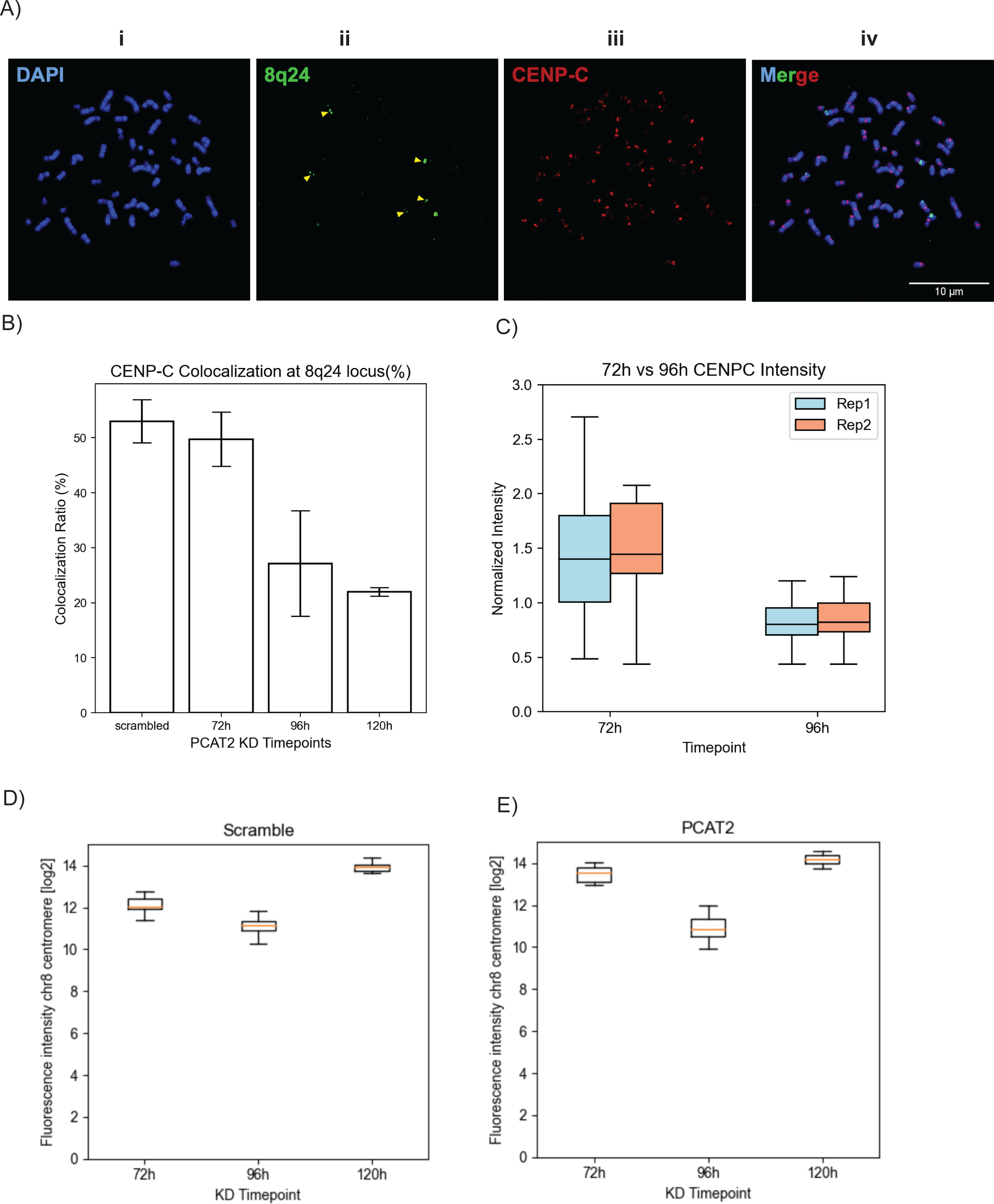
MetaChrome quantification reevaluates CENP-C intensity and colocalization at the ectopic 8q24 locus and native centromeres. (A) Representative immunofluorescence-FISH (IF-FISH) image of a metaphase spread from SW480 cells. Chromosomes are stained with DAPI (blue, i), the 8q24 locus is identified by FISH probe (green, ii), and CENP-C protein is detected by immunofluorescence (red, iii). The merged image (iv) shows colocalization. Scale bar = 10 µm. MetaChrome quantification of CENP-C at the ectopic 8q24 locus. (B) CENP-C colocalization ratio (%) with the 8q24 locus following PCAT2 knockdown (KD) at 72, 96, and 120 hours, compared to a scramble control (Error bars represent standard deviation, n = 2). (C) Normalized mean CENP-C fluorescence intensity (relative to scramble control) at the 8q24 locus for replicate 1 and replicate 2 at 72- and 96-hours post-transfection. (D) MetaChrome quantification of CENP-C fluorescence intensity (log2 scale) at the native chromosome 8 centromere in SW480 cells following knockdown of PCAT2. (E) compared to a scramble control at 72 hours, 96 hours, and 120 hours post-transfection. Representative multi-FOV images collected at three timepoints (72 h, 96 h, and 120 h) for scramble controls, with sample counts of (30, 30, 11) and (18, 32, 21) fields of view, respectively.

## 4. Discussion

Here we describe MetaChrome, a new biological image software platform that automates and simplifies the analysis of metaphase chromosomes. To our knowledge, MetaChrome is the first open-source software that helps researchers automate the analysis of images of mitotic chromosomes taken in a fluorescence microscope. Because it is open-source and modular, MetaChrome’s core segmentation can be further trained or adapted by other users for their specific experimental questions, for example, by retraining the segmentation model with user-annotated datasets. MetaChrome also addresses bottlenecks in high-throughput image analyses for this task by automating labor-intensive chromosome segmentation and quantification of IF and/or DNA-FISH signals, thereby reducing analysis time and human error.

The major challenge in the automated analysis of images of mitotic chromosomes is their object instance segmentation. To address this, we have fine-tuned an existing CellPose model [25] to optimize it for mitotic chromosome segmentation. The fine-tuned Cellpose model demonstrated significantly improved performance in correctly separating mitotic chromosomes (F1 score >0.8 at 0.7 IoU, Fig. 3G), representing an advancement over standard automated approaches. Our comparative analysis highlighted the inherent weaknesses of conventional watershed-based segmentation, which frequently struggled with clustered chromosomes and led to substantial errors (Fig. 3C). Even the “naïve” Cellpose model exhibited limitations, underscoring the necessity of dataset-specific fine-tuning to achieve reliable segmentation for chromosomes in fluorescence microscopy images. For this work, the fine-tuning was conducted on a limited dataset (26 images before augmentation and 1835 chromosomes-see Materials and Methods for technical details). Recognizing the diverse needs of researchers, we include straightforward instructions for additional fine-tuning of the Cellpose model described in the Cellpose 2.0 paper [25] using new, user-annotated metaphase chromosome datasets. These custom-trained models can then be easily integrated into the existing workflow, allowing users to leverage all of MetaChrome’s features on datasets for which it was not originally trained.

MetaChrome’s modular software architecture, built on scientific open-source software libraries and integrated into the versatile Napari visualization platform, expands its accessibility and utility to a broader scientific community. This graphical user interface enables users to load and visually inspect images, set image analysis parameters on a few representative images, and then run the analysis in batch mode on larger datasets. To further enhance usability with novel datasets and accommodate the inevitable biological variability inherent in metaphase chromosome samples, we have incorporated an option for manual refinement of segmentations, enabling users to correct instances where the automated process may not have perfectly identified chromosome boundaries. Importantly, these manually corrected segmentations can be saved and subsequently used as training masks, facilitating a continuous cycle of improvement for future model retraining efforts. This flexibility, combining automated power with manual oversight, is necessary for precise downstream quantifications, particularly in fluorescence in situ hybridization and colocalization studies, where accuracy in spot detection directly influences biological conclusions. Furthermore, our automated detection of DNA-FISH and protein signals, complemented by these manual refinement capabilities, ensures reliable identification of relevant chromosomal loci. Altogether, our findings clearly show the advantages of using a specially retrained deep-learning model within an intuitive graphical interface, improving accuracy, reproducibility, and throughput.

To validate MetaChrome capabilities, we used MetaChrome to quantify CENP-C fluorescence intensity at both the 8q24 ectopic site and chromosome 8 centromere following the knockdown of specific 8q24-derived lncRNAs, such as PCAT2 [28]. Our software accurately measured the decrease in CENP-C intensity specifically at the 8q24 locus upon PCAT2 knockdown, while confirming the stability of CENP-C levels at the native centromere under the same conditions. These results precisely replicate the findings reported previously [28], showcasing MetaChrome’s power not only for discovery but also for rigorously validating published data regarding site-specific protein localization changes. By offering robust and reproducible segmentation results, as exemplified by the validation of lncRNA knockdown effects on CENP-C localization, MetaChrome facilitates accurate comparisons across experimental conditions and between research groups, laying the foundation for standardized and comparable chromosome analyses across different experimental settings.

MetaChrome has certain limitations when it comes to segmenting chromosomes from aged cells and specific cell types that exhibit metaphase chromosomes in fuzzy and loosely packed structures. Notably, if metaphase chromosomes are digested in pepsin for an extended period, the chromatin can become detached from the compact metaphase structure, posing challenges for the segmentation process [42–44]. However, this issue can be addressed by training the model with manually curated images. Additionally, when two different targeted DNA-FISH or protein colocalization spots are located close to each other on a specific chromosome, there may be an overlap in the area considered for intensity measurement. This complication can be mitigated through demultiplexing the analysis.

To enhance MetaChrome utility, future efforts will focus on expanding and diversifying annotated chromosome datasets, particularly for challenging samples, and integrating the novel Cellpose-SAM model [45] to improve object segmentation. Additionally, we plan to incorporate features like automated multi-channel colocalization analysis from two different DNA-FISH signals from different target regions to broaden MetaChrome’s applicability to basic research and clinical uses. Furthermore, we aim to advance MetaChrome’s capabilities in measuring chromosome length, detecting translocations or deletions, identifying marker chromosomes, and recognizing chromothripsis [46,47]. Finally, we aim to integrate and overlay data acquired from diverse imaging technologies such as atomic force microscopy (AFM), high-resolution images from super-resolution microscopy, X-ray diffraction, and cryo-electron microscopy (cryo-EM) [48–54] to study protein colocalizations, target gene locus, DNA breaks, deletions, and translocations in greater detail. By pursuing these initiatives, we aim to establish MetaChrome as a more powerful and versatile tool for chromosome analysis.

In conclusion, MetaChrome represents a significant advancement in automated metaphase chromosome analysis, successfully integrating state-of-the-art deep learning methods, user-friendly visualization, and quantitative analytical capabilities validated against existing biological research. Future improvements could focus on expanding annotated datasets, further refining deep-learning models with various annotated chromosome datasets, and integrating additional analytical functionalities to broaden its utility across diverse experimental and clinical contexts.

## 6. Acknowledgements

This work utilized the computational resources of the NIH HPC Biowulf cluster (https://hpc.nih.gov). We thank Prabhakar Reddy Gudla for key initial contributions on image analysis of mitotic chromosomes that eventually led to MetaChrome. We would also like to thank Sweta Sikder and Annie Gilbert for their assistance in testing the software and providing valuable feedback. Finally, we would like to thank Truman McNeil and Tom Misteli for their critical reading of the manuscript.

This research was supported by the Intramural Research Program of the National Institutes of Health (NIH) (Projects 1-ZIC-BC011567-11 and 1-ZIA-BC011206-16). The contributions of the NIH author(s) were made as part of their official duties as NIH federal employees, are in compliance with agency policy requirements, and are considered works of the United States Government. However, the findings and conclusions presented in this paper are those of the author(s) and do not necessarily reflect the views of the NIH or the U.S. Department of Health and Human Services.

